# Passive neuromodulation: an energy-driven mechanism for closed-loop suppression of epileptic seizures

**DOI:** 10.64898/2026.03.26.714592

**Authors:** Gagan Acharya, Andrew Huang, Vijayalakshmi Santhakumar, Erfan Nozari

## Abstract

For decades, electrical neuromodulation has been used as a therapeutic mechanism to disrupt and desynchronize pathological neural activity in various neurological disorders. Despite notable progress, however, patient outcomes remain highly variable, particularly in medically intractable epilepsy where surgery still provides the greatest chance of seizure freedom. Here we propose *passive neuromodulation (PNM)* as a radical alternative to conventional neurostimulation, whereby analogue feedback is used to drain energy from an epileptic circuit and thus suppress the initiation or spread of electrographic seizures. We provide pilot evidence on the efficacy and robustness of PNM using two computational models of epileptic dynamics: a detailed biophysical network model of dentate gyrus, and the Epileptor neural mass model of seizure dynamics. Despite the vast differences between these models, our results show the robust ability of PNM to suppress seizures in both models. We further demonstrate the efficacy and robustness of *responsive PNM*, whereby brief (50ms) windows of PNM are triggered by a simultaneously-running seizure detection algorithm, as well as the safe and tunable nature of PNM, where more robust seizure suppression can be achieved by parametrically titrating the amount of power drained from the tissue, without inducing any seizures even if applied interictally. Overall, our results provide strong evidence on the promise of PNM for the closed-loop control of epileptic seizures and other neurological disorders where damping pathological network activity can restore healthy dynamics.

## Introduction

About 3 million people in the United States and 50 million worldwide have active epilepsy, with about 30% having drug-resistant epilepsy (DRE)^7, 11, 83^. Yet, few treatment options with significant risks and side effects are clinically available for DRE. The greatest chance of seizure freedom comes from surgical resection–one of the most invasive treatments in medicine^47, 63^. Yet, 20-25% of patients who undergo invasive pre-surgical intracranial EEG (iEEG) monitoring do not qualify for surgery^71^, and among the patients who undergo surgery only 60-70% become seizure-free^78, 88^.

The costs and risks of resection have long motivated seizure suppression via neuromodulation, both open-loop^21, 59^ and closed-loop^5, 6, 54^, with the latter leading to the development of the Responsive Neu-rostimulation (RNS) system^35, 72^. However, despite significant improvements in seizure frequency—over 50% seizure reduction in about 73% of patients implanted by the Responsive Neurostimulation (RNS) system^55^, e.g.—the majority of patients implanted by all existing neuromodulation systems continue to have seizures, with some even experiencing an increase in their seizure frequency^55, 67, 81^. This is ex-acerbated by a great amount of preoperative unpredictability in individual responses and the months-to-years needed before reaching maximum efficacy, even in responders^55, 67^.

Several studies have sought to design and test more effective alternatives in animals^5, 20, 40, 45, 66, 74, 80^ as well as human patients^12, 32, 50, 87^. Despite notable progress, however, no existing work has been able to achieve complete seizure freedom in all or even a predictable subset of subjects using electrical neurostimulation. A critical need, therefore, exists to develop robust, predictable, and mechanistically grounded algorithms for neuromodulation in epilepsy. The present study takes a radical step towards this goal by revisiting the fundamentals of neuromodulation and how it affects seizure dynamics from an energy perspective.

For over half a century, seizures have been viewed to emerge from an imbalance of excitation and inhibition, leading to an unstable feedback loop and run-away excitation^30, 38, 52^. This view aligns well with the standard definition of epileptic seizures as ‘abnormal excessive or synchronous neuronal activity’^22^ and has persisted as the dominant view and the basis for most theoretical models of seizure generation in epilepsy^23, 37, 61^. Models based on bistability, e.g., are a notable extension of the classical view that provides a more nuanced explanation of what happens when interictal activity loses its stability^17, 23, 37, 48^. Despite their differences, however, nearly all existing theories of ictogenesis attribute the transition from normal towards seizure activity to a progressive loss of stability through destabilizing, positive feedback.

Unstable dynamics, regardless of their precise form or mechanism, lead to massive production of energy^41^. In a seizure, this energy manifests in different forms, including electromagnetic power embedded in the seizure’s paroxysmal electrographic fluctuations, which then feed back to the neuronal circuit via ephaptic coupling^52, 69^. Historically, different approaches have been taken to break this cycle, including more advanced metabolism-targeting^25, 27, 51, 64^ as well as pure thermal treatments^73, 76, 85^. These approaches have proven quite successful, and thus support the premise of targeting energetics, but they rely on metabolic or thermodynamic processes that are inherently slow.

In this work, we provide pilot evidence that unstable seizure energetics can be robustly controlled using near-instantaneous electrical damping, an approach we refer to as passive neuromodulation (PNM). PNM is supported by half a century of theoretical and empirical research on passivity-based control in the engineering literature^42, 79, 82^. A system is called *passive* if it dissipates energy and *active* otherwise. Therefore, passive systems are naturally stable, and passivity-based control functions by adding passive elements, such as Ohmic resistors in an electrical circuit or friction-based dampers in mechanical systems, to drain the excess energy of an unstable system. This approach has proven successful in various electromechanical systems^14, 15, 70^ but, to our knowledge, has never been used or tested in neuroengineering. The present study aims to translate passivity-based control for the closed-loop control of epileptic seizures and provide pilot evidence on its efficacy using in silico computational models of seizure dynamics.

## 1 Results

### Passive neuromodulation: the basic principle

The core principle of PNM is to ensure that the neuromodulator circuit is passive, i.e., that it can only drain energy from the underlying tissue (Figure 1a). To achieve this in an electrically active tissue such as the brain, consider a neuromodulator that reads the differential local field potential Δ*LFP* (*t*) between two microelectrodes and, in response, passes an electrical current *I*_*P NM*_ (*t*) through the tissue via the same pair of electrodes (Figure 1b). It then follows from basic circuit theory (the definitions of electrical potential and current)^2^ that the product

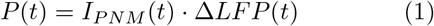

equals the instantaneous power that is transferred *from* the neuromodulator *to* the underlying tissue.^∗^ If Δ*LFP* (*t*) is in *µ*V and *I*_*P NM*_ (*t*) in mA, *P* (*t*) will be in nW, and the integral 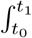 gives the total energy in nJ that is transferred *from* the neuromodulator *to* the underlying tissue between times *t*_0_ and *t*_1_.

**Figure 1:**
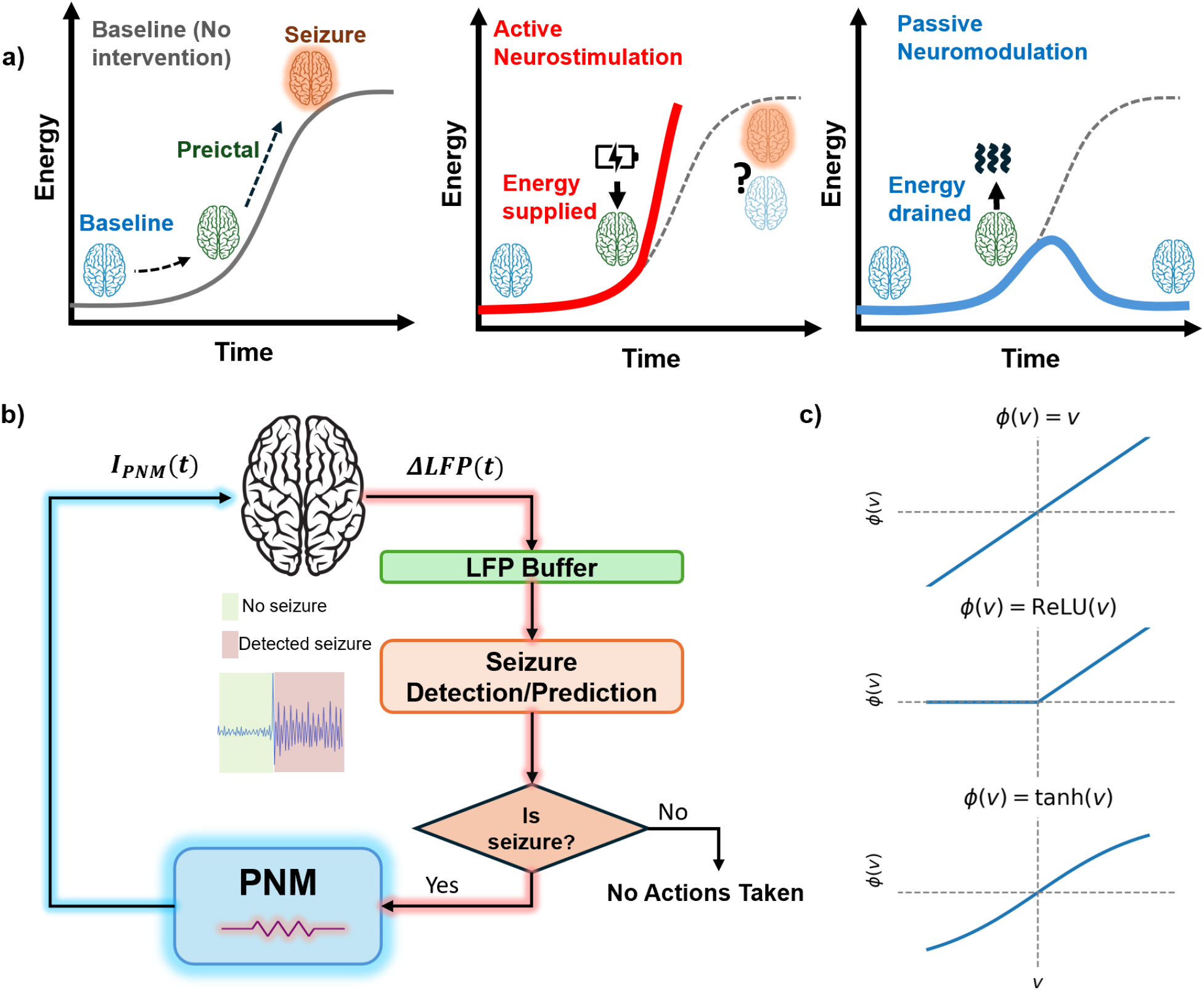
Overview of Passive Neuromodulation (PNM). **(a)** Schematic showing the difference between active and passive neuromodulation. Left panel shows the energy landscape of the interictal to ictal transition in the absence of interventions. Throughout this work, ‘energy’ refers to electromagnetic energy (such as that recorded by all forms of EEG) unless otherwise stated. The effect of active neurostimulation (middle panel) on this landscape can vary, fairly unpredictably, from seizure suppression to even seizure induction depending on the parameters of stimulation and the neural populations recruited (cf. Figure 2). In contrast, PNM (right panel) monitors and adjusts the amount and direction of the electrical current passed through the tissue such that energy is always drained from it. Thus, the result is “cooling down” the tissue and bringing the network back to its baseline (interictal) state. **(b)** High-level block diagram of PNM together with concurrent seizure detection or prediction. The differential local field potential Δ*LFP* across (at least one) pairs of microelectrodes are recorded and monitored over a moving window (buffer) for seizure detection/prediction. Upon detection/prediction, PNM applies the current *I*_*P NM*_ through the same pair of microelectrodes for a brief period of time (typically *<*1s) with polarity such that energy is solely drained from the tissue (cf. Eq. (3)). **(c)** Representative examples of conductance profiles *ϕ* satisfying the passivity condition in Eq. Eq. (4). Shown are a linear (Ohmic) profile, a ReLU-type gated profile, and a tanh profile. All satisfy *V ϕ*(*V* ) ≥ 0, thereby guaranteeing that energy is dissipated rather than injected.

Accordingly, the neuromodulator is called passive if

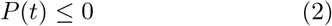

for all time *t*. This can be guaranteed if the neuro-modulator applies the current

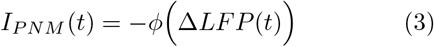

where *ϕ*, called the ‘conductance profile’, is a function that satisfies

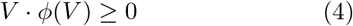

for all *V*. There are many such functions *ϕ*, some of which are shown in Figure 1c. The simplest choice is the linear function *ϕ*(*V* ) = *g*_*P NM*_ *V* corresponding to an Ohmic resistor with conductance *g*_*P NM*_ *>* 0. It is straightforward to show (and quite intuitive) that if the underlying tissue is itself passive, then its connection with any passive neuromodulator (any function *ϕ* satisfying Eq. (4)) remains passive and therefore stable^42^.

However, when the underlying tissue is electrically active, especially when it is intrinsically unstable, such as epileptic tissue, interconnecting it with a passive neuromodulator can only stabilize the tissue if the neuromodulator is sufficiently ‘strong’, i.e., the energy drained by the neuromodulator exceeds the energy generated by the tissue. This not only depends on the conductance profile *ϕ*, but also on its placement in space (electrode locations) and time (seizure detection). These three factors thus constitute the key design variables of any PNM, including the ones studied in this work. It is important to note, however, that even a poorly designed PNM is at worst ineffective–unlike an active stimulator, a passive controller can never exacerbate instability (induce seizures) because of its passive nature.

### PNM effectively suppresses mesial temporal lobe seizures when placed at the seizure onset zone (SOZ)

We first validated the ability of PNM to suppress epileptic seizures in a biophysical model of the dentate gyrus (DG), a critical node in the generation and spread of mesial temporal lobe (MTL) seizures^9, 18, 34, 46, 58^. The model includes over 500 multi-compartmental excitatory granule and mossy cells as well as inhibitory basket cells and hilar interneurons with axonal projections to the perforant-path termination zone. When implemented with recurrent mossy fiber sprouting, the model exhibits prolonged seizure-like activity after focal stimulation of the perforant path inputs (Figure 2a-b, see Ref.^68^ for details). PNM was applied via Eq. (3) with

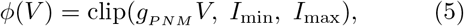

with *g*_*P NM*_ = 50S, *I*_min_ = 0, and *I*_max_ = 2mA (Figure 2a). The PNM microelectrodes were placed at the SOZ between the granule cell (GC) dendritic trees (Figure 2a) and the neuromodulator was turned on for 100ms following seizure onset (Methods).

**Figure 2:**
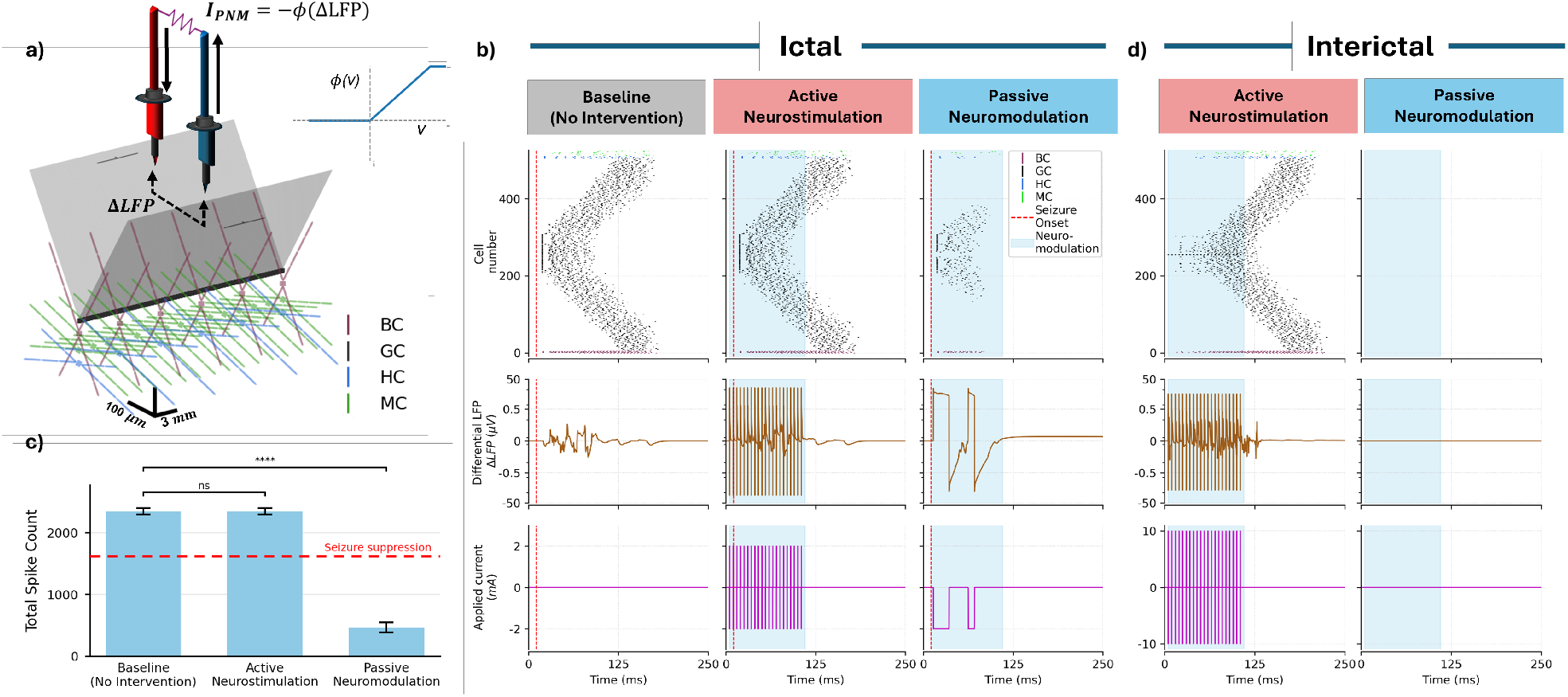
Reliable seizure suppression with single-site PNM at the seizure onset zone (SOZ). **(a)** 3D schematic of the dentate gyrus (DG) biophysical NEURON model used to simulate seizure-like activity and its response to PNM (see Methods). A pair of microelectrodes delivers brief PNM at the midpoint location between the granular cell (GC) dendritic trees. In the baseline model, seizures originate from the middle 100 GCs following perforant path input, constituting the SOZ. **(b)** Raster plots of single-unit spiking activity (top row), differential local field potential between the pair of microelectrodes (Δ*LFP*, middle row), and current injected/drained via the same pair of microelectrodes (bottom row) under baseline, active neurostimulation (250 Hz, 2 mA), and PNM conditions. Active neurostimulation has no significant impact, while PNM reliably stops the seizure. **(c)** Distribution of total number of single-unit spikes under each condition in (b). Bar plots show mean ± 1 s.e.m. PNM achieves *>* 80% reduction in total number of spikes (*p <* 0.01, Wilcoxon signed-rank test), which is well below the threshold for successful seizure suppression (see Methods for calculation of suppression threshold). Active neurostimulation leads to no significant change in the total number of spikes. **(d)** Effects of active neurostimulation and PNM in the absence of baseline seizures. Active neurostimulation can induce a seizure if applied with sufficiently large amplitude, whereas PNM leads to no visible perturbations of the underlying network.

PNM achieved reliable and near-complete suppression of seizure activity, with over 4-fold reduction in mean total spike count (*p <* 0.01, Wilcoxon signed-rank test) (Figure 2b,c). In contrast, active neurostimulation resulted in no significant change in seizure activity, allowing it to continue, spread, and terminate as it would without intervention (*p >* 0.95, Wilcoxon signed-rank test). Furthermore, unlike active neurostimulation that can provoke a seizure if applied interictally (Figure 2d, see also^16, 48, 57^), the passive nature of PNM prevents it from exciting the tissue even if turned on during the interictal state. Therefore, PNM is not only significantly more effective in seizure suppression, but it is also inherently safe by design.

### Multi-site PNM can reliably suppress seizures with arbitrary onset zones

We next examined the dependence of PNM efficacy on its proximity to the SOZ. To do so, we modified the DG model such that its SOZ can be parametrically varied along the GC axis (see Methods), and simultaneously varied the location of a single pair of PNM contacts independently parallel to the same axis (Figure 3a). We measured PNM efficacy via

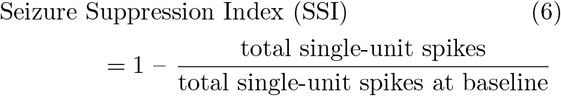

and monitored its dependence on the location of the SOZ and its distance from the PNM contacts. Here, we focus on granule cell spikes, as they constitute the dominant population activity and are the principal excitatory neurons in the dentate gyrus. As shown in Figure 3b, the efficacy of single-site PNM is fairly insensitive to the location of the SOZ and mainly dependent on the PNM-SOZ distance. Seizure suppression remains near baseline (SSI ≃ 0) when PNM contacts are 20mm or farther from the SOZ, and monotonically improves with reduced PNM-SOZ distance (Figure 3c). At the PNM-SOZ distances of 2mm or smaller, single-site PNM reaches its maximum efficacy, with an average SSI of 64% across all seizure onset zones (Figure 3d). Therefore, even with a single pair of electrode contacts, PNM has the potential to immediately and effectively suppress seizures– significantly better than active neurostimulation–in cases such as focal structural epilepsies where the SOZ is small and accurately identifiable via imaging.

**Figure 3:**
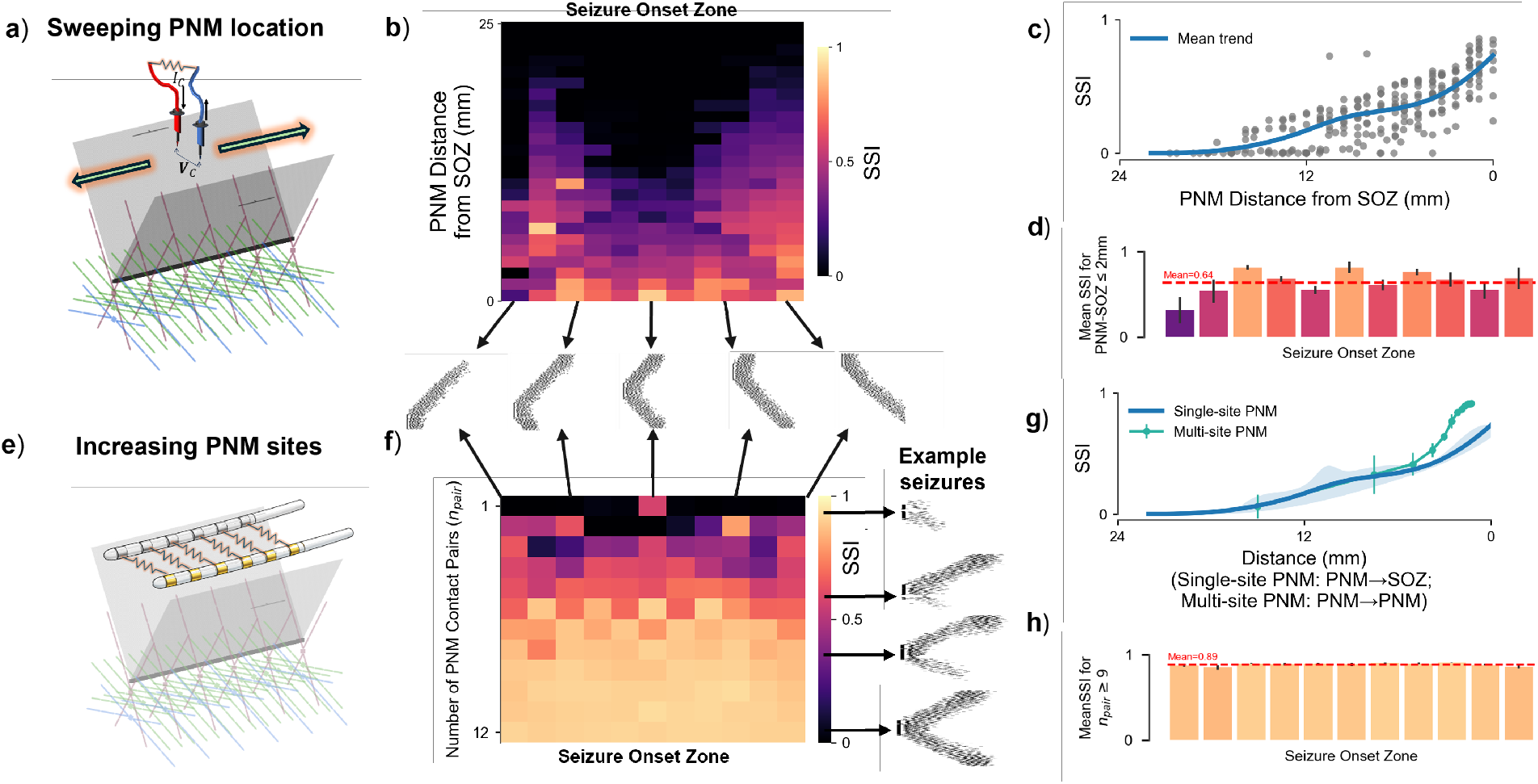
Relaxing the need for SOZ targeting with multi-site PNM. **(a)** Schematic diagram of the same DG model as in Figure 2 but with the SOZ and the location of a single PNM site parametrically swept along the GC axis (see Methods). **(b)** Heatmap of Seizure Suppression Index (SSI, Eq. (6)) as a function of SOZ location and the horizontal distance between the PNM microelectrodes and the SOZ. Representative rasters below the heatmap illustrate how the seizure spreads from different SOZs. Seizure suppression is strongest when PNM is near the SOZ and decays with the PNM-SOZ distance. **(c)** Scatter plot of SSI as a function of PNM-SOZ distance. Each dot represents one cell in (b) and the blue line represents the smoothed average. Suppression depends on and improves monotonically with decreasing distance between PNM contacts and the SOZ. **(d)** Distribution of mean SSIs from (b) when PNM contacts are within 2mm of the SOZ. Error bars show 1 s.e.m. Suppression is effective across all SOZ locations, with a grand mean SSI of over 64%. **(e)** Schematic diagram of multi-site PNM in the same DG model. All contacts lie on a single pair of electrodes placed longitudinally along the main DG axis. **(f)** Similar heatmap as in (b), but with the vertical axis showing the number of PNM contact pairs. Representative rasters on the right illustrate sample seizures corresponding to different SSIs. Seizure suppression is near-complete with 9 or more contact pairs, regardless of the location of the SOZ. **(g)** Mean SSI of single- and multi-site PNM as a function of the PNM-SOZ distance (single-site PNM, blue) or distance between PNM contact pairs (multi-site PNM, green). The blue line is the same as that in (c). Error shades and bars represent 2 s.e.m. The two trends are mostly consistent, with multi-site PNM becoming significantly more effective in smaller distances. **(h)** Distribution of mean SSIs ± 1 s.e.m. from (f) with 9 or more PNM contact pairs. Suppression is uniform and near complete, with a grand mean SSI of 89%.

We next studied the case where the SOZ cannot be accurately mapped, and increased the number of PNM contact pairs to maintain seizure suppression efficacy. To do so, we rotated the orientation of the microelectrodes to lie longitudinally parallel to the main DG axis, and placed up to 12 equally-spaced contacts on each electrode (Figure 3e). The SOZ was varied along the GC axis as before, while the PNM contacts remained fixed and uniformly covering the length of the DG (extreme case of no SOZ information/random SOZ). Seizure suppression improved significantly with increasing the number of PNM contact pairs (Figure 3f,g), reaching a maximum efficacy of 89% with nine or more pairs (2 ∼ 3mm-apart contact spacing, Figure 3). This is remarkable, in particular, given the worst-case nature of this experiment, where the seizures could originate from any SOZ across the DG network, and demonstrate the potential of multi-site PNM for robust real-time seizure suppression even in the absence of accurate SOZ localization and/or the presence of multiple seizure foci.

### Multi-site PNM is robust to delays in PNM onset

Moving towards a fully closed-loop neuro-modulation scheme where both the timing and applied current of PNM are decided in real time (Figure 1), we next analyzed whether PNM remains effective if it is turned on with some delay, allowing the seizure to spread and intensify before PNM gets a chance to stop it. Despite the clear advantages of proactive neuromodulation^20, 80^ and the possibility of forecasting seizures far in advance significantly better than chance^1^, accurate seizure prediction remains an open problem, and closed-loop neuromodulation remains largely reactive. As such, the robustness of any closed-loop neuromodulation scheme to its onset latency remains a critical factor in determining its practical utility. We analyzed this robustness by implementing single- and multi-site PNM similar to above, but now activating PNM with variable amounts of delay relative to seizure onset time (Figure 4).

**Figure 4:**
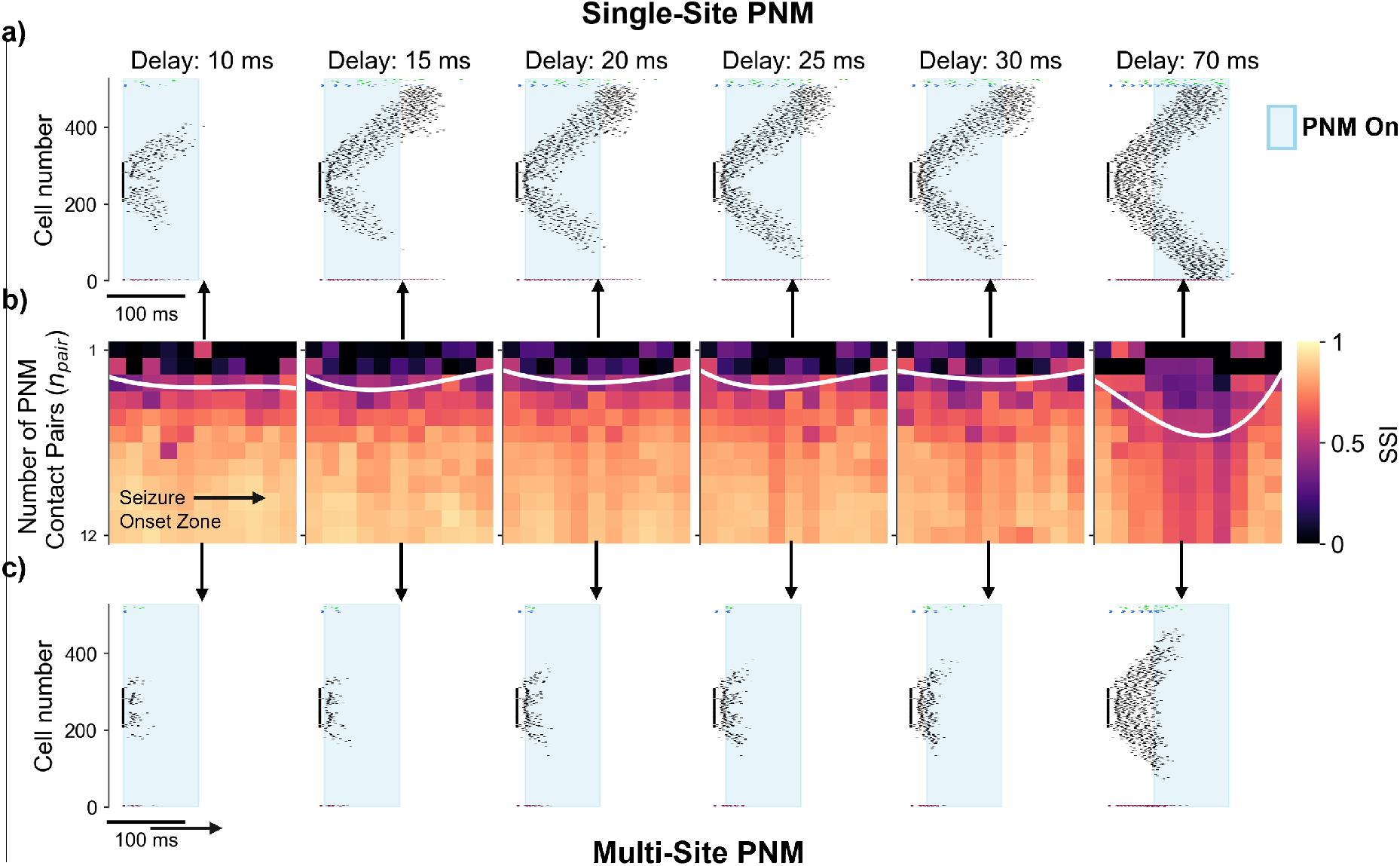
Robustness of PNM to delays in PNM onset. **(a)** Raster plots of single-unit activity under 100 ms of single-site PNM with increasing onset delays. SOZ and the PNM contact pair are kept at the center while PNM onset delays were increased (delays shown with respect to the time of seizure onset, see Methods). Single-site suppression is robust to about 10ms of delay, which is while seizure spread remains within the spatial reach of the single pair of electrodes. Suppression gradually loses efficacy under longer delays and becomes ineffective at about 70ms delay. **(b)** Heatmap of Seizure Suppression Index (SSI, Eq. (6)) as a function of SOZ location, number of multi-site PNM contact pairs, and varying PNM onset delays. The layout of each heatmap is similar to Fig 3f. The white line shows an approximate threshold for successful seizure suppression (see Methods). Suppression efficacy remains nearly unchanged up to about 30ms delay, and highly robust even at 70ms delay. Note that SSI calculation is based on all single-unit spikes (in the numerator), including those that occurred before PNM onset–hence the darker shades in the last heatmap. **(c)** Similar to (a) but for multi-site PNM with 12 contact pairs. Seizures are immediately suppressed post-PNM onset, regardless of the PNM latency.

Our results demonstrate a strong robustness of PNM to onset delays, particularly with 7 or more PNM contact pairs (Figure 4b). Single-site PNM is only weakly robust to onset delays, maintaining its original efficacy up to about 10ms of delay (Figure 4a). Note, importantly, that the limiting factor is not the passage of time per se, but the spatial spread of seizure activity beyond the radius of effectiveness of the single pair of PNM contacts (as seen before in Figure 3a-d). As such, increasing the number of PNM contact pairs and the spatial coverage that ensues from it significantly improves robustness to onset delays. With 12 contact pairs (2.5 mm intercontact spacing), PNM remains effective and near-instantaneously suppresses an ongoing seizure, even if it is turned on after the seizure has recruited over half of the network (Figure 4c). Thus, PNM efficacy depends largely on its spatial reach, and remains highly robust to delays in seizure detection as long as seizure spread remains within the radius of effectiveness of the available array of PNM contact pairs.

### Responsive PNM triggered by simultaneous seizure detection

Supported by the robustness of multi-site PNM to uncertainties in the location (Figure 3e-h) and time (Figure 4c) of seizure onset, we tested the ability of multi-site PNM to suppress spontaneously-generated seizures in a fully closedloop setting with concurrent seizure detection in the loop. We modified the deterministic DG model described so far so that seizures can emerge spontaneously at random times and locations as a result of background stochastic input to the network (Methods). At baseline, the resulting model generates a variety of seizure events, including partial and multifocal seizures (Figure 5c).

**Figure 5:**
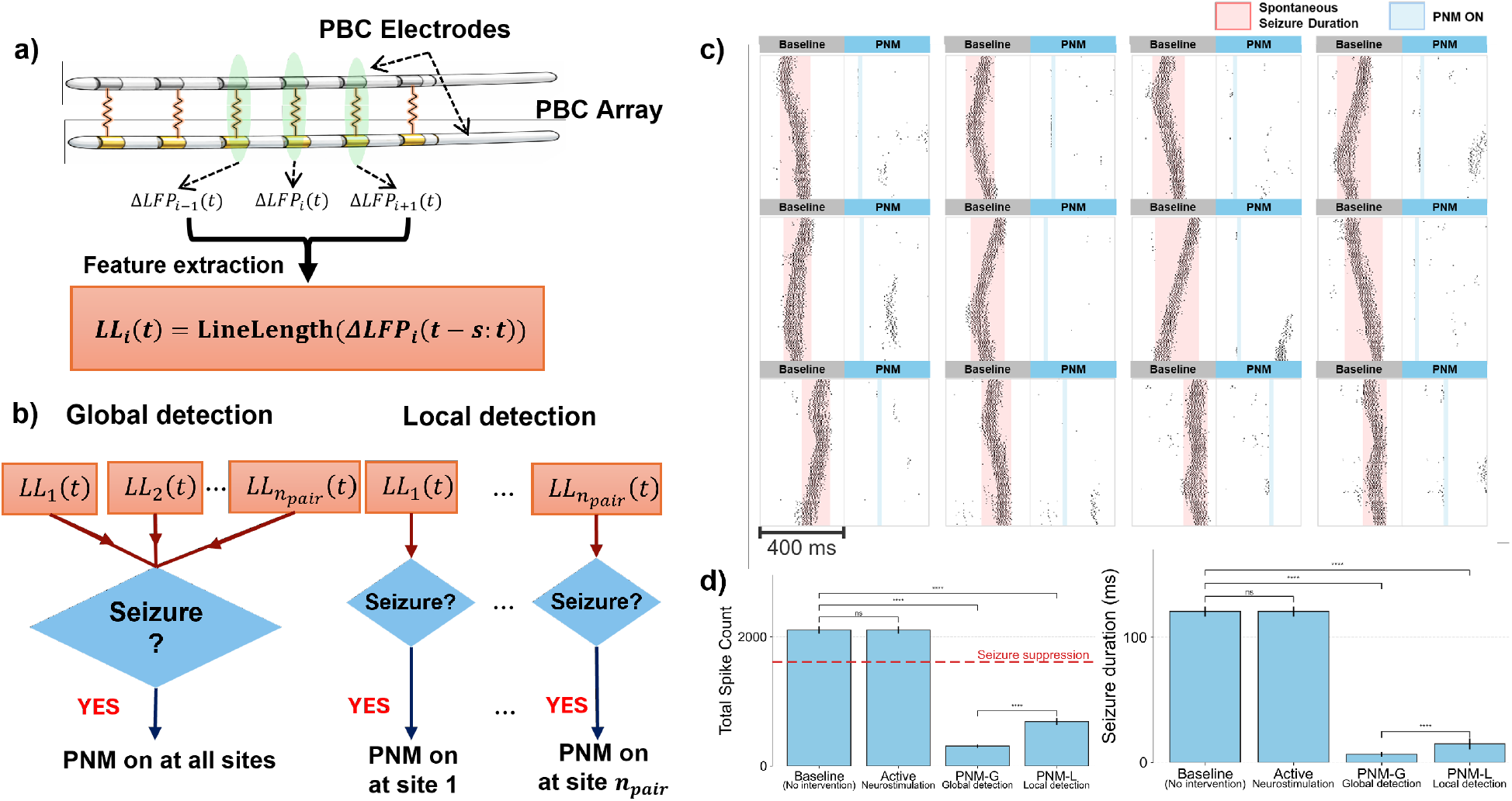
Suppression of spontaneous seizures with simultaneous seizure detection and PNM. **(a)** Schematic of feature extraction used for seizure detection. The differential local field potential Δ*LFP*_*i*_ between each pair of PNM contacts (cf. Eq. (3)) over a moving window of *s* = 10ms was used to compute line length for each pair *i* (see Methods). **(b)** Seizure detection was then performed using random forest classifiers either globally (using all pairs) or locally (using each pair separately). A detection in the global setting triggered a synchronous 50ms PNM event across all contact pairs, whereas each detection in the local setting triggered an asynchronous 50ms PNM event across the respective pair only. **(c)** Sample scatter plots of single-unit spiking activity in the spontaneously-seizing DG model with and without detection-triggered PNM. All panels show the results of global detection followed by synchronous PNM across 11 contact pairs. Each pair of Baseline-PNM panels are identical (use the same random seed) except for the PNM application. Seizures are effectively and near-instantaneously detected and suppressed in all cases. **(d)** Mean total single-unit spike count (left) and seizure duration (right) under baseline, active neurostimulation, and PNM with global and local seizure detection. Seizure duration is estimated from the onset and offset times of population spiking activity, identified using the same thresholding criteria used in seizure labeling for the training of seizure detection classifiers (Methods). Error bars show 1 s.e.m., and the dashed horizontal line shows the threshold for successful seizure suppression (see Methods). Both forms of detection-triggered PNM were able to successfully suppress all seizures, with global detection leading to faster suppression at the cost of larger overall intervention (greater power drained from the tissue).

To detect the onset of spontaneous seizures in the stochastic model, we designed a simple seizure detection algorithm based on the line length of the same differential LFPs used for computing PNM current (Figure 5a, see Methods for details). Compared to complex algorithms we have developed before for seizure forecasting^1^, the present algorithm, mimicking the simplest detection criteria used by the RNS system^75^, is computationally efficient to ensure feasibility on a chronic implant. We tested the performance of two detection paradigms (Figure 5b). In a ‘global’ paradigm, seizure events were detected based on the line length features across all PNM contact pairs. Upon each detection, PNM was correspondingly activated across all contact pairs for a limited duration of 50ms. In a ‘local’ (a.k.a. distributed) paradigm, seizure detection was performed independently for each contact pair, and 50ms of PNM was applied across only the respective contact pair upon each detection. Compared to the former, the local paradigm is ‘minimal’ in that it applies significantly less overall current in a spatially targeted and refined manner.

Given the stochastic nature of the model, we performed *n* = 1000 trials with and without PNM, and compared the seizure suppression efficacy of local and global detection-triggered PNM against baseline and active neurostimulation. Figure 5c shows several illustrative examples of single-unit spiking activity at baseline vs. global detection-triggered PNM. PNM effectively suppresses seizure activity in nearly all trials, regardless of the time, location, or nature of seizure onset. Figure 5d quantifies these effects, showing the total number of single unit spikes and seizure duration across baseline, active neurostimulation, and local and global detection-triggered PNM. Similar to deterministic time-triggered seizures (Figure 2), active neurostimulation has no significant effect on seizure intensity or duration, whereas both PNM paradigms damp network activity in a reliable and reproducible manner. Notably, while the global activation paradigm is significantly more effective—owing to the simultaneous engagement of all PNM pairs and the resulting broader spatial extent of energy dissipation—the local activation paradigm, in which only pairs associated with detected seizure activity are activated, is also sufficient by a wide margin to achieve successful suppression. Overall, these results provide pilot evidence on the efficacy of fully closed-loop PNM, whereby the reactive application of PNM is sufficient to suppress seizures that have already spread broadly enough to be detectable in the line length of the local field potential.

### PNM in a phenomenological neural mass model of seizure dynamics

To test the limits of PNM and better understand its impact on seizure dynamics, we further implemented PNM in the Epileptor model of Jirsa and colleagues^37^ (Figure 6a). The epileptor is a low-dimensional neural mass model widely used to simulate seizure-like activity, both as a lumped model and as a distributed network^36, 48, 60, 65^ (see Methods). We applied PNM to this model following the same dissipative feedback form in Eq. (3) with the conductance profile

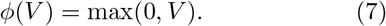

**Figure 6:**
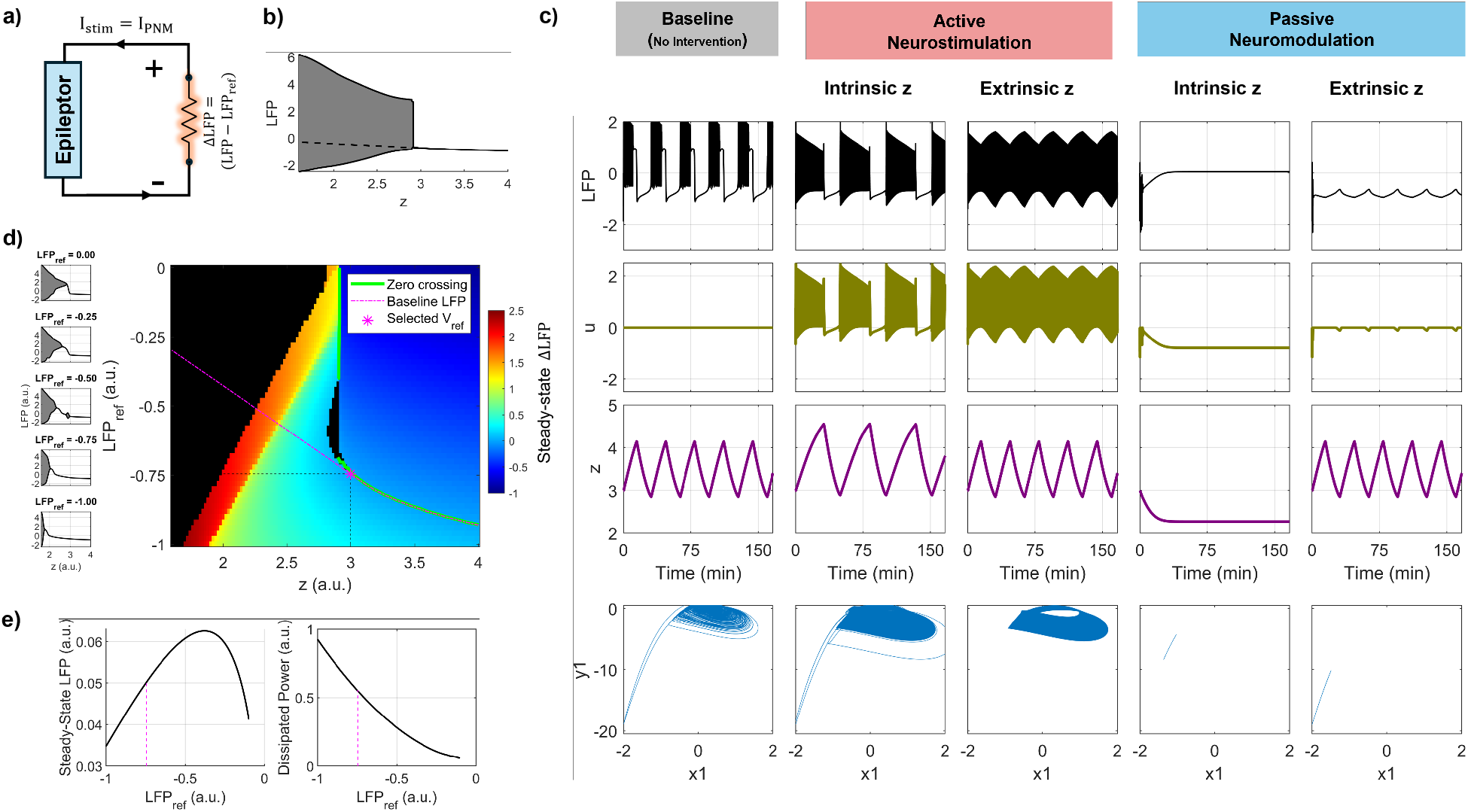
Passive neuromodulation of the Epileptor. Epileptor is a phenomenological neural mass model widely used to simulate seizure-like activity in an abstract and theoretically well-understood manner. **(a)** Schematic of the closed-loop electrical circuit formed by interconnecting the epileptor with PNM. Even though the epileptor is comprised of normative differential equations over abstract variables, we treat its two input variables (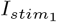 and 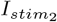) as modeling electrical currents and its output variable (*x*_2_ − *x*_1_) as modeling the LFP. The differential LFP, Δ*LFP* = *LFP* − *LFP*_*ref*_ ), is used for feedback, where *LFP*_*ref*_ denotes the interictal (equilibrium) value of *x*_2_ − *x*_1_. **(b)** The bifurcation diagram of the epileptor model at baseline (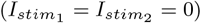). *z* denotes the slow permittivity variable that acts as the bifurcation parameter. Pathological oscillations reminiscent of ictal activity occur for *z* ≲ 3 (shaded area, denoting the range of oscillations) whereas a stable equilibrium modeling interictal activity is reached for *z* ≳ 3 (solid line). The latter was used as *LFP*_*ref*_ and extrapolated, when needed, to the ictal regime (dashed line). **(c)** Detailed comparisons of epileptor dynamics at baseline as well as under active and passive neuromodulation. Both forms of neuromodulation were tested on the original model (‘Intrinsic *z*’, where the slow permittivity variable followed its differential equation in Eq. (11)) as well as under an ‘Extrinsic *z*’ condition where *z* was forced to follow its baseline trajectory. Under both conditions, active neurostimulation exacerbates the seizures, whereas PNM completely suppresses them. **(d)** The heatmap of steady-state (equilibrium) differential LFP (*LFP* − *LFP*_*ref*_ ) under PNM as a function of different values of *z* (kept fixed, as in (b)) and *LFP*_*ref*_. Cases where no equilibrium was reached (seizure activity/oscillatory steady-state persisting in spite of PNM) are shown in black. The green line (zero crossing) separates the region where the model is by-itself seizure-free (*I*_*P NM*_ = 0), and the region where seizures are suppressed by the PNM (*I*_*P NM*_ *<* 0). Note that by increasing *LFP*_*ref*_, the model can tolerate smaller *z* values without falling into a seizure. **(e)** Impact of *LFP*_*ref*_ on the steady-state LFP and dissipated power (power drained by the PNM) when *z* follows its intrinsic dynamics. Note the increased power dissipation (increased PNM engagement) as *LFP*_*ref*_ is increased, corresponding to the bottom (most seizure resistant) side of the heatmap in (d).

As we will see below, the half-wave rectification in Eq. (7) is critical for turning on PNM only when needed for seizure suppression. We set Δ*LFP* equal to the difference between the model’s LFP output (i.e., *x*_1_ − *x*_2_, see Methods) and its output’s steadystate equilibrium value *LFP*_*ref*_ ,

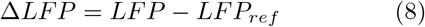

and applied the resulting *I*_*P NM*_ to the model’s *I*_*stim*_ (cf. Eq. (11)). The subtraction of *LFP*_*ref*_ is equivalent to a change of coordinates and accounts for the fact that, unlike the DG model, the ‘reference’ (interictal) LFP in the epileptor is nonzero. Given the well-understood dependence of ictogenesis in this model on the slow permittivity variable *z* (Figure 6b), and the unknown possible effects of PNM on such variables in the brain (extracellular potassium, oxygen, etc.)^37^, we applied PNM under two conditions. Under the ‘Intrinsic *z*’ condition, *z* remains endogenously generated (following Eq. (11)) and, consequently, affected by the PNM. Under the ‘Extrinsic *z*’ condition, in contrast, *z* was exogenously forced to follow its time course at baseline, independent of the effects of PNM on other variables.

Under both intrinsic-*z* and extrinsic-*z* conditions, PNM was able to fully suppress seizures, while active neurostimulation exacerbated them (Figure 6c). Note that, at baseline, the slow permittivity variable *z* follows a sawtooth trajectory oscillating at about 30Hz between approximately 2.9 and 4.1 (a.u.). Under intrinsic-*z* active neurostimulation, this oscillation slows down to about 20Hz (prolonged seizures), and *z* needs to reach a higher value (about 4.5, more interictal) for a seizure to stop. If *z* is forced to follow its baseline trajectory (Extrinsic *z*), seizures never stop under active neurostimulation. In contrast, under PNM with intrinsic *z*, neuromodulation is able to completely stop seizures, *even though z* reacts to PNM by settling at a much smaller (more ictogenic) value of about 2.3. Under the extrinsic-*z* condition the job of PNM is ‘easier’, preventing seizures with minimal intervention (*I*_*P NM*_ ≠ 0) applied only when *z* falls too low (≲ 3).

Figure 6d further demonstrates the robustness of epileptor seizure suppression via PNM and the critical role played by the reference output *LFP*_*ref*_ in PNM’s robustness. As *LFP*_*ref*_ is increased, so does the range of *z* values for which PNM can suppress seizures (colored area). In other words, for any value of *LFP*_*ref*_, there exists a sufficiently small value of *z* where PNM fails, but increasing *LFP*_*ref*_ pushes this threshold on *z* to lower values and, accordingly, increases the robustness of seizure suppression via PNM. Mechanistically, this increased robustness stems from an increased engagement of PNM. Note that, from a controls perspective, Eq. (8) denotes the tracking error which the application of *I*_*P NM*_ seeks to bring to zero. Increasing *LFP*_*ref*_ makes this error more negative, thus increasing *I*_*P NM*_ and the amount of power that PNM drains from the epileptor (Figure 6e). Therefore, *LFP*_*ref*_ acts as a control knob that one can use to balance the tradeoff between achieving more robust seizure suppression and doing so with less exogenous intervention.

## 2 Discussion

### Summary

In this study, we provided pilot evidence on the efficacy of passive neuromodulation (PNM) as a radical approach to closed-loop control of seizures in drug-resistant epilepsy. In contrast to the status quo, we used neuromodulation via passive analogue circuitry to damp, rather than excite, the surrounding tissue. Using a biophysical model of dentate gyrus exhibiting seizure-like activity as baseline, we provided strong evidence on the seizure suppressive efficacy and robustness of PNM, provided that seizure spread remains within the spatial reach of PNM—approximately 2mm. We guaranteed the latter via multi-site PNM, and demonstrated its near-complete seizure suppressive efficacy when combined with real-time seizure detection. Despite major differences in spatiotemporal scale and level of abstraction, PNM was also able to completely suppress seizures in the Epileptor neural mass model of seizure dynamics. The simplicity of this model allowed us to further analyze PNM as an error-driven feedback control mechanism (similar to a voltage clamp) and demonstrate the strong relationship between the robustness of seizure suppression and the amount of power drained by the PNM.

### Related work

Several studies have explored closed-loop electrical stimulation for control of epileptic seizures. The Responsive Neurostimulation system^35, 72^ was originally designed as a real-time closed-loop system that delivers bursts of neurostimulation in order to terminate detected electrographic seizures^54^. Mounting evidence suggests, however, that its efficacy in patients who respond to it stems mainly from long-term network reorganizations (plasticity) rather than real-time seizure suppression^44, 62^. The idea of real-time closed-loop seizure suppression has nevertheless remained popular, with several attempts in animals^5, 20, 40, 45, 66, 74, 80^ and human patients^12, 32, 50, 87^. To our knowledge, the closest attempt to PNM is that of Gluckman and colleagues^29^ who implemented a proportional (P) controller in hippocampal slice preparations, showing that immediate seizure suppression could be achieved and maintained for several minutes. While this approach demonstrated robustness in a controlled setting, it mapped power to voltage rather than voltage to current, leading to periods of tissue excitation during interictal phases and rebound seizures upon deactivation. Subsequent modifications, such as half-wave rectification, mitigated some of these issues but still introduced charge imbalance and variable effects. These limitations highlight the need for a control strategy that is inherently passive, stable, and avoids unintended excitation.

Our work is also related to and builds on the rich literature on the computational modeling of seizure dynamics. Biophysically detailed network models have been widely used to study seizure initiation, propagation, and control^31, 49, 77, 86^, providing a valuable substrate for the pilot testing of neuromodulation strategies. Low-dimensional models such as the Epileptor, on the other hand, have also been extensively studied as a way to capture the essential dynamics underlying seizure transitions in a reductionistic framework^3, 8, 13, 19, 26, 36, 39, 53, 84^. Our work builds on this literature and uses models from both scales of analysis to test the strengths and limitations of PNM.

### PNM Design and Implementation

Our pilot results clearly demonstrate that, while PNM is generally safe due to its passive nature, its ability to suppress seizures relies on its proper design and implementation. Our experiments with the DG model show that both the proximity of the PNM contact pairs to the SOZ and their spatial orientation (dipole vector direction) play an important role in PNM efficacy. In particular, we found PNM to be most effective when the PNM contacts were placed near the GC dendritic trees with their anode-cathode dipole oriented, asymmetrically, perpendicular to the dendritic planes. Likewise, we expect the identification of proper electrode configuration and conductance profile to constitute an important design stage in any application of PNM. In our experience, however, the range of electrode placements and conductance profiles that led to clearly recognizable efficacy was large enough to be easily discoverable without much trial and error.

Compared to a linear (Ohmic) conductance profile *ϕ*(*V* ) = *g*_*P NM*_ *V*, we used a saturated and half-wave rectified current (Eq. (5)) to ensure safety and improve the stability of the applied current. We empirically observed that for a given bound on the amplitude of injected current, steep conductance profiles (*g*_*P NM*_ ≫ 1 with immediate saturation) are more effective than shallow profiles (*g*_*P NM*_ ≪ 1 with graceful saturation). The former is reminiscent to the so-called ‘bang-bang control’ that is known to be optimal for minimum-time control problems^43^. Likewise, in our experiments, we found half-wave rectified profiles to result in more stable current trajectories with less chattering than full-wave profiles. In other words, restricting currents to negative (or positive) values reduces oscillatory behavior (rapid attempts to self-correct) in the feedback loop and yields more reliable suppression of pathological activity.

While beyond the scope of this work, we anticipate subsequent real-world implementations of PNM to involve digital circuitry for seizure monitoring and detection and analogue circuits for control. The latter, while being in principle feasible with completely passive elements, may require an active ‘rendering’ of passive behavior depending on the complexity and scales of the desired conductance profiles. Regardless of design procedures, however, PNM holds significant promise as a framework for closed-loop control of epileptic seizures owing primarily to its conceptual simplicity and transparent mechanism.

### Limitations

Our findings of PNM efficacy and characterization are limited to the two computational models we have used as baseline; the biophysical DG model and the Epileptor. Future work is needed to replicate these findings experimentally, starting with animal models, for which the present study provides significant insights. Moreover, given the focus of our baseline models on the dynamics of seizures, our experiments cannot directly address the potential effects of PNM on other, non-seizure-related functions of the underlying circuits. In particular, the time-invariant nature of our models limits our ability to study potential interactions between PNM and neural plasticity. We anticipate, however, that PNM’s impact on non-seizure dynamics will be generally minimal, particularly in responsive PNM, compared to active neurostimulation due to the passive and minimally disruptive nature of PNM. From a theoretical perspective, a further limitation of this work is the lack of analytical guarantees for closed-loop stability under PNM. As noted, PNM builds on a rich history of passivity-based control in engineering, where analytical guarantees are often pursued as safeguards against potentially overlooked failure modes. While beyond the scope of this work, we foresee analytical characterizations to be feasible for the Epileptor model owing to its low-dimensional structure, constituting valuable avenue for future research. High-dimensional biophysical models such as that used in this work, however, often do not lend themselves to more than numerical simulations, as conducted herein.

### Conclusions

We proposed passive neuromodulation (PNM) as a contrastive alternative to all existing forms of neuromodulation for the control of focal seizures in drug-resistant epilepsy. We provided proof-of-concept demonstrations of PNM in two computational models of epileptic seizures at different scales, and paved the way for subsequent experimental testing of PNM. If successful, PNM can provide a transformative paradigm shift in closed-loop neuromodulation for epilepsy and beyond based on the robust and mechanistically transparent principle of passivity.

## 3 Methods

### Biophysical Dentate Gyrus Model

To evaluate the efficacy of PNM for seizure suppression, we employed as baseline a biophysical computational model of the dentate gyrus (DG). The model, described before in^68^, is a simplified yet anatomically faithful network scaled approximately 2000:1 from the rodent DG. It consists of 527 multi-compartment neurons simulated in the NEURON environment^10, 33^, including 500 granule cells, 15 mossy cells, 6 basket cells, and 6 hilar perforant-path–associated (HIPP) interneurons. The neurons are arranged within a lamellar topology that preserves the principal excitatory and inhibitory motifs of the DG circuitry. The model reproduces seizure-like discharges resulting from pathological mossy fiber sprouting and cell loss and was thus used as a physiologically grounded baseline for evaluating seizure-suppression strategies.

We used an extended version of this model proposed in^28^ whereby the cells were spatially embedded along a one-dimensional anatomical axis (Figure 2a). This spatial representation is critical for realistic computation of extracellular transfer resistances (depending on the distances between cell segments and the neuromodulation site) which are needed to properly calculate local field potentials (LFPs) as well as neuromodulation-evoked neuronal responses. In the original NeuroConstruct implementation^28^, neurons were positioned along a 1,000 µm axis corresponding to a rodent DG slice. To approximate the scale of the human DG, we uniformly scaled this spatial extent by a factor of 30, resulting in an effective DG length of approximately 30 mm. The original connectivity and synaptic architecture were nevertheless preserved.

### Simulation Protocol

All simulations involving the DG model were conducted in the NEURON environment with a fixed integration time step of Δ*t* = 0.02 ms. Each simulation was run for a total duration of *T* = 250 ms in deterministic experiments (i.e., seizures initiated by perforant path input) and *T* = 400 ms for stochastic simulations with spontaneous seizures. The somatic membrane potentials of all 527 neurons as well as the LFPs and extra-cellular currents were continuously recorded. Action potentials were detected using cell-type–specific voltage thresholds, consistent with the original model^68^: GCs: −48.7 mV, MCs: −52 mV, BCs: −49 mV, and HCs: −50 mV.

### Incorporation of Stochastic Background Activity

The original model of^68^ consists of deterministic dynamics, i.e., the same input always generates the same response. To simulate physiological variability and ongoing synaptic drive, we augmented this model by adding stochastic background inputs to a random subset of neurons. To maintain biophysical realism, background activity was locally structured via input groups, namely, compartments that were spatially close along the neuron morphology were assigned to the same group, resulting in locally correlated fluctuations, while maintaining statistical independence across distant regions. The size of each group was variable, allowing for natural heterogeneity in the spatial extent of correlated inputs, with a mean group size controlled by a hyperparameter *g*_max_. In experiments requiring deterministic seizure initiation—consistent with the original model^68^—the parameter *g*_max_ was set to a relatively low value (approximately 10) to preserve predictable seizure onset and prevent spontaneous activity. In contrast, experiments evaluating responsive PNM with concurrent seizure detection employed a higher value of *g*_max_ (approximately 50) to facilitate spontaneous seizure generation.

Background input was applied in the form of persistent spike trains with a mean rate of approximately 20 Hz. Each spike produced an exponentially decaying synaptic current, with amplitude randomly drawn between 0 and 0.05 nA to introduce moderate variability in input strength. Approximately 45% of neurons received background inputs, ensuring that network activity remained heterogeneous and realistic. Granule cells were assigned slightly higher input amplitudes, between 0 and 0.1 nA, to reflect summated afferent/perforant path inputs.

### Seizure Initiation via Perforant Path Input

In all experiments except those involving simultaneous seizure detection, seizure-like activity was elicited through activation of the perforant path (PP) afferents to DG as in the original model^68^. A brief external current pulse (1 ms duration, 0.5 nA amplitude) was applied at a predefined time (t = 10 ms) via the PP input channel. This stimulation reliably evokes seizure-like discharges beginning approximately 20 ms after pulse onset. The PP input was connected to the dendritic compartments of the middle 100 GCs (out of 500 total) and the middle 2 BCs (out of 6 total), corresponding to a focal seizure onset zone (SOZ) near the middle of the 30 mm longitudinal extent of the model.

To parametrically vary the location of SOZ along the DG axis, we shifted the indices of the granule and basket cells receiving PP input as

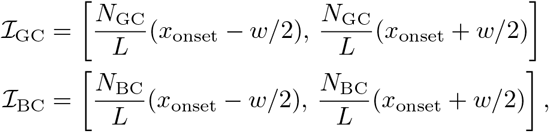

where *x*_onset_ ∈ [0, *L*] is the desired SOZ location, *L* = 30 mm is total DG length, *N*_GC_ = 500 and *N*_BC_ = 6 denote the total number of granule and basket cells, respectively, and *w* = 6 mm is the spatial width of the initiation region.

### Spontaneous Seizure Initiation

Spontaneous seizures were induced by disabling the perforant path input and increasing the degree of background correlation in the network. This was achieved by using a high grouping parameter (*g*_max_ ≃ 50, see above), leading to broader spatial correlations in synaptic noise. This enhanced the likelihood of localized clusters of hyperactivity emerging spontaneously within the network, which in turn initiated seizure-like discharges at random locations and times without perforant input.

### Modeling Extracellular Fields and Currents (Neuromodulation)

We considered a bipolar electrode configuration for recording differential LFPs and applying extracellular current. Let the anode and cathode locations be at **p**_*a*_ = (*x*_*a*_, *y*_*a*_, *z*_*a*_) and **p**_*c*_ = (*x*_*c*_, *y*_*c*_, *z*_*c*_), respectively. Then, for each cellular compartment at position **r** = (*x, y, z*), the *extracellular transfer resistance* can be approximated assuming a heterogeneous and isotropic medium with constant resistivity *ρ*,

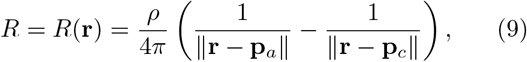

where ∥ · ∥ denotes the Euclidean distance. We used a resistivity of *ρ* = 400 Ω · cm, corresponding to reported low-frequency human gray matter conductivities of 0.2–0.3 S*/*m^4, 24^. The extracellular potential induced at each cellular compartment by an extracellularly-applied (active or passive) current *I* can then be computed via Ohm’s law:

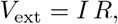

*V*_ext_ is then applied to each compartment using the NEURON’s `extracellular` mechanism (`xtra.mod` and `extracellular.mod`), allowing compartments to experience the computed extracellular potential while preserving the original intracellular dynamics. For simulations involving multiple bipolar contacts, the extracellular potential induced at the location of any compartment was computed as a linear superposition of those induced by each contact pair, and vice versa.

The same mechanism, applied in reverse, can be used to calculate the differential LFP induced between each electrode contact pair by the currents flowing in cellular compartments. At each time step, the contribution of a given compartment to Δ*LFP* was estimated as the product of its transmembrane current density and the effective extracellular transfer resistance between the compartment and the contact pair. The results were then summed over all neuronal compartments, giving

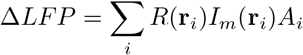

where *I*_*m*_(**r**_*i*_) is the transmembrane current density in the *i*^*th*^ compartment, *A*_*i*_ is the corresponding membrane area, and *R*(**r**_*i*_) is the transfer resistance in Eq. (9).

### Electrode Placement

Bipolar electrode pairs (anode–cathode) were oriented perpendicular to the granule cell longitudinal axis. The anode and cathode contacts were separated by 100 *µ*m, a distance determined empirically through separate hyperparameter tuning experiments. Each pair was positioned symmetrically with respect to and placed 300 *µ*m above the GC axis.

For deterministic experiments involving seizure initiation via PP input and a single electrode pair, the optimal axial placement depended on the seizure onset zone. Specifically, the electrode pair was centered at the longitudinal location corresponding to the center of the granule cell group receiving PP input, as defined in the seizure initiation protocol.

For experiments involving multiple electrode pairs (both spontaneous and PP-driven seizures), axial placement was performed symmetrically along the GC axis. Neighboring electrode pairs were separated by a uniform distance of *L/n*_pair_, where *n*_pair_ is the number of electrode pairs.

### Seizure Suppression Threshold Estimation

To quantify how much a neuromodulation strategy must reduce spiking—beyond natural trial-to-trial variability—to be considered effective, we computed a delay-dependent seizure suppression threshold from baseline spontaneous-seizure simulations (different random seeds). Only trials with exactly one detected seizure were retained.

For each retained trial, we computed the total number of granule cell spikes occurring after seizure onset, with an additional delay *δ* ∈ {0, 10, 15, 20, 25, 30, 70 } ms. This yielded, for each delay, a set of baseline spike counts

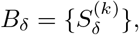

where 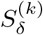 denotes the total post-onset spike count in trial *k* at delay *δ*. To reduce sensitivity to rare low-count realizations, we discarded the lowest 10% of values in *B*_*δ*_. We then estimated intrinsic multiplicative variability by repeatedly sampling random pairs *b*_*i*_, *b*_*j*_ ∈ *B*_*δ*_ (50,000 samples per delay) and computing the log-ratio

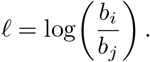

This procedure characterizes the distribution of relative spike-count differences expected purely from stochastic baseline fluctuations.

The suppression threshold at delay *δ* was then defined as

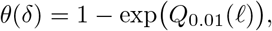

where *Q*_0.01_(*ℓ*) denotes the 1st percentile of the logratio distribution. Consequently, *θ*(*δ*) represents the minimum fractional reduction in post-onset spiking that exceeds 99% of baseline variability. This delay-dependent threshold was used to declare successful seizure suppression in subsequent analyses.

### Seizure Detection and Control Activation

In experiments involving responsive PNM, seizure detection was implemented using a random forest (RF) classifier. The line length of each differential LFP trace Δ*LFP*_*i*_ was computed over a sliding 10 ms window (equivalent to *s* = 500 samples) as

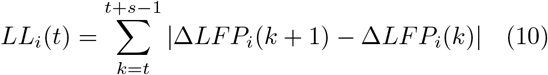

and provided to the RF classifier for detection. The classifiers were trained (with 5-fold cross-validation) using data from spontaneous seizure simulations conducted without any control intervention. A total of 200 trials were generated, each exhibiting varying seizure onset times and spatial locations. For labeling, the simulation time was divided into non-overlapping 5 ms bins, and each bin was assigned a *seizure* label if the spike count across all GCs exceeded a threshold corresponding to 5% of the total GC population. To avoid transient false positives, a bin was labeled as a seizure only if this condition persisted for all bins within the subsequent 20 ms interval. This criterion ensured robust labeling of genuine seizure onsets while maintaining temporal precision by detecting the earliest onset point.

As noted above, two detection strategies were evaluated. Under *global detection*, a single classifier receives the concatenated line-length features of all PNM sites and makes a single decision as to whether seizure activity is present anywhere within the network. A global detection was followed by 50ms of PNM application across all contact pairs. In contrast, under *local detection*, independent RF classifiers monitor the differential LFP line length at each site and trigger the local application of 50ms PNM at the respective site upon seizure detection. The latter approach minimizes the application of PNM not only over time but also over space.

### Active Stimulation

PNM was compared against an active stimulation paradigm. Inspired by the responsive neurostimulation (RNS) system^56^, the stimulation consisted biphasic 160 *µ*s current pulses at 200 Hz. The length, timing, and peak current of active stimulation events were kept the same as those of PNM to facilitate side-by-side comparisons.

### Experiments Involving the Epileptor Model

The Epileptor model^37^ is defined by a system of six coupled ordinary differential equations, herein simulated as

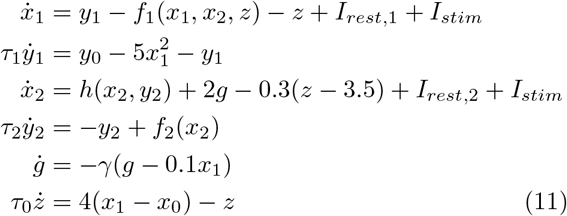

where 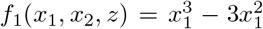 if *x*_1_ *<* 0 and (*x*_2_ − 0.6(*z* −4)^2^)*x*_1_ otherwise, *f*_2_(*x*_2_) = 6(*x*_2_ +0.25) if *x*_2_ ≥ −0.25 and 0 otherwise, and 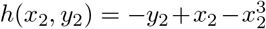. Following the original model^37^ we used

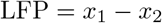

as the LFP output used for closed-loop neuromodulation.

#### Passive and active neuromodulation

PNM was applied to the epileptor as described in Results (cf. Eq. (7) and Eq. (8)). Active neurostimulation was applied similarly, except using the conductance profile *ϕ*(*V* ) = − *V* which gives *V* · *ϕ*(*V* ) ≤ 0 (opposite to Eq. (4), violating the passivity condition).

#### Simulation protocol

Simulations of the Epileptor model were performed using a fixed-step Euler integration of Eq. (11). Key simulation parameters are summarized in Table 1.

**Table 1:** Epileptor simulation parameters.

| Parameter | Value |
| --- | --- |
| Integration step size, $\Delta t$ | 0.01 s |
| Simulation duration, $t$ | 0–6000 s |
| Initial conditions, $y(0)$ | $[0, -5, z_{\text{init}}, 0, 0, 0]^T$ |
| Slow permittivity variable range, $z$ | 4.0 to 1.6, step $-0.005$ |
| Fast subsystem offset, $x_0$ | $-1.6$ |
| Fast subsystem baseline, $y_0$ | 1 |
| Fast time constant, $\tau_1$ | 1 |
| Slow time constant, $\tau_0$ | 2857 |
| Slow subsystem time constant, $\tau_2$ | 10 |
| Resting input, $I_{rest,1}$ | 3.1 |
| Resting input, $I_{rest,2}$ | 0.45 |
| Permittivity damping, $\gamma$ | 0.01 |

#### Bifurcation diagrams

The bifurcation diagrams shown in Figure 6 were generated by systematically varying the slow permittivity variable *z* and recording the resulting LFP extrema. Maximum and minimum LFP values were plotted against *z*, with partial Gaussian smoothing applied for *z <* 2.95 to better illustrate the transition boundaries. Under PNM, neuromodulation shifts the bifurcation boundaries, reducing the LFP range over which seizure-like activity occurs.

#### Evaluation conditions

As noted in Results, we considered two evaluation scenarios based on the dynamics of the slow permittivity variable *z*. In the absence of any control (*I*_*stim*_ = 0), *z* exhibits a sawtooth oscillation, ranging between *z*_min_ ≈ 2.9 and *z*_max_ ≈ 4.0. Under the ‘intrinsic-z’ condition, we allowed *z* to be generated intrinsically by the model, thus remaining coupled with other state variables according to its original differential equation (Eq. (11)) and, thus, becoming indirectly affected by neuromodulation. In contrast, under the ‘extrinsic-z’ condition, we provided *z* as an **extrinsic** variable to the model, forcing it to follow the same sawtooth trajectory as it did at baseline.

## Acknowledgments

We would like to sincerely thank Dr. Ted Carnevale for his invaluable guidance on simulating PNM using the NEURON environment. This work was supported in part by the National Science Foundation under award #2239654 to E.N., by the National Institutes of Health (NIH) NINDS grants R37NS069861 and R01NS097750 to V.S., and by NIH/NINDS F31NS131052 and AES award 957615 to A.H.

## Author Contributions

E.N. designed and supervised the study; G.A. performed the analyses; A.H. and V.S. provided guidance on the biophysical dentate gyrus model; all authors contributed to writing the manuscript.

## Code availability

The code used to generate the results in this study is available at https://github.com/nozarilab/2026Acharya_PNM

## Footnotes

∗ This is assuming the standard sign convention where *I*_*P NM*_ leaves the cathode/enters the anode of the neuromodulator and Δ*LFP* = *LFP*_*cathode*_ − *LFP*_*anode*_.

